# Cell Autonomous Neuroprotection by the Mitochondrial Uncoupling Protein 2 in a Mouse Model of Glaucoma

**DOI:** 10.1101/522821

**Authors:** Daniel T. Hass, Colin J. Barnstable

## Abstract

Glaucoma is a group of disorders associated with retinal ganglion cell (RGC) degeneration and death. There is a clear contribution of mitochondrial dysfunction and oxidative stress toward glaucomatous RGC death. Mitochondrial uncoupling protein 2 (*Ucp2*) is a well-known regulator of oxidative stress that increases cell survival in acute models of oxidative damage. The impact of *Ucp2* on cell survival during sub-acute and chronic neurodegenerative conditions however is not yet clear. Herein, we test the hypothesis that increased *Ucp2* expression will improve retinal ganglion cell survival in a mouse model of glaucoma. We show that increasing retinal ganglion cell but not glial *Ucp2* expression in transgenic animals decreases glaucomatous RGC death, but also that the PPAR-γ agonist rosiglitazone, an endogenous transcriptional activator of *Ucp2*, does not significantly alter RGC loss during glaucoma. Together, these data support a model whereby increased *Ucp2* expression mediates neuroprotection during a long-term oxidative stressor, but that transcriptional activation alone is insufficient to elicit a neuroprotective effect, motivating further research in to the post-transcriptional regulation of *Ucp2*.

## Introduction

Glaucoma is a group of disorders associated with retinal ganglion cell (RGC) degeneration and death (Quigley, 2011) and, after cataracts, is the most frequent cause of blindness worldwide (Resnikoff and Keys, 2012). An increase in intra-ocular pressure (IOP) is a prominent risk factor for glaucoma (Boland and Quigley, 2007), and most therapeutic solutions designed to prevent RGC death in glaucoma share the ability to decrease IOP. However, IOP reduction does not reduce glaucomatous visual field loss in roughly half of patients (Leske et al., 2004), necessitating development of adjuvant therapeutic modalities, including neuroprotective molecules that can protect RGCs.

The molecular mechanisms of glaucoma pathogenesis are multifactorial, but are frequently connected to an increase in damaging free radicals in the eye (Izzotti et al., 2003; Saccà and Izzotti, 2008), retina (Tezel et al., 2005), and optic nerve head (Malone and Hernandez, 2007; Feilchenfeld et al., 2008), herein termed oxidative stress. Ocular hypertension increases RGC oxidative stress (Chidlow et al., 2017), despite anti-oxidative support from endogenous antioxidant proteins (Munemasa et al., 2009) and resident glial cells of the retina, including astrocytes and müller glia (Varela and Hernandez, 1997; Carter-Dawson et al., 1998; Kawasaki et al., 2000; Woldemussie et al., 2004). Exogenous addition of serum antioxidants such as vitamin E are not necessarily protective from the disease (Ramdas et al., 2018), suggesting that current anti-oxidative therapeutics for glaucoma are insufficient.

Mitochondria are a well-known source of cellular free radicals, which during oxidative phosphorylation can leak from multi-protein complexes of the electron transport chain such as NADH:Ubiquinone Oxioreductase and Coenzyme Q:Cytochrome C Oxioreductase (St-Pierre et al., 2002). Mitochondrial ROS production is greater at hyperpolarized mitochondrial membrane potentials (Ψ_m_), and in isolated mitochondria small decreases in Ψ_m_ significantly decrease levels of ROS (Korshunov et al., 1997; Miwa et al., 2003). Endogenous uncoupling proteins, particularly the mitochondrial uncoupling protein 2 (UCP2) are able to protect nervous tissue from multiple sources of acute damage (Mattiasson et al., 2003; Andrews et al., 2005; Lapp et al., 2014; Barnstable et al., 2016) by decreasing Ψ_m_ and presumably ROS (Fleury et al., 1997; Echtay et al., 2001). Lower levels of ROS are protective in most scenarios, but the predicted outcome of a lower Ψ_m_ is also a decreased mitochondrial drive for ATP synthesis(Klingenberg and Rottenberg, 1977). Therefore it is unclear whether uncoupling proteins are beneficial for long-term neurodegenerative conditions. As with many neurodegenerative disorders, the clinical course of glaucoma progresses over multiple years. It is therefore essential that model systems of neurodegeneration develop over time and not in reaction to a single damaging insult.

In the microbead model of glaucoma, occlusion of the irido-corneal angle progressively increases damage to RGCs over a sub-acute time frame (Huang et al., 2018). Using this model we tested whether enhanced *Ucp2* expression in mouse RGCs or in supporting glial cells is protective against injury. We found that increasing levels of *Ucp2* in RGCs, but not in GFAP-expressing glia, was neuroprotective. *Ucp2* levels are under several forms of transcriptional and translational control (Donadelli et al., 2014; Lapp et al., 2014), and our second goal was to determine whether factors that increase *Ucp2* transcription provide protection from cell death. We found that the PPAR-γ agonist rosiglitazone, a well known transcriptional activator of Ucp2, does not alter RGC survival during glaucoma, implying an additional need to characterize clinically useful molecules which regulate *Ucp2* at post-transcriptional levels.

## Materials and Methods

### Ethical approval

This study was carried out in accordance with the National Research Council’s Guide for the Care and Use of Laboratory Animals (8th edition). The protocol was approved by the Pennsylvania State University College of Medicine Institutional Animal Care and Use Committee.

### Animals

Wild-type C57BL6/J (WT) and transgenic mice were housed in a room with an ambient temperature of 25°C, 30-70% humidity, a 12-hr light–dark cycle, and ad libitum access to rodent chow. Transgenic mouse strains, B6.Cg-Tg(*GFAP-cre/ER*^*T2*^)505Fmv/J (*Gfap-creER*^*T2*^, Stock#: 012849) (Ganat et al., 2006) and Tg(*Thy1-cre/ER*^*T2*^,-EYFP)HGfng/PyngJ *(Thy1-creER*^*T2*^, Stock#: 012708) (Young et al., 2008) were each generated on a WT background and purchased from the Jackson Laboratory (Bar Harbor, ME, USA). *GFAP-creER*^*T2*^ and *Thy1-creER*^*T2*^ mice express a fusion product of *cre* recombinase and an estrogen receptor regulatory subunit (*creER*^*T2*^) under the control of the *hGFAP* or *Thy1* promoters, respectively. CreER^T2^ activity is regulated by the estrogen receptor modulator and tamoxifen metabolite 4-hydroxytamoxifen (Zhang et al., 1996). Ucp2KI^fl/fl^ mice were derived from Ucp2KOKI^fl/fl^ mice (provided by Sabrina Diano, PhD) and result from multiple back-crosses with WT mice (Toda et al., 2016). In these crosses, mice were selectively bred to retain the Ucp2KI sequence and the WT variant of the *Ucp2* gene. In these mice, a transgene was inserted in to the R26 locus, containing a LoxP-flanked stop codon followed by a copy of the mouse *Ucp2* cDNA and an IRES-EGFP sequence. Following cell-type specific cre-mediated excision of the LoxP-flanked stop codon, these mice express *Ucp2* and EGFP in astrocytes and müller glia (*Ucp2KI*^*fl/fl*^; *GFAP-creER*^*T2*^ mice) or in the vast majority of retinal ganglion cells (*Ucp2KI*^*fl/fl*^; *Thy1-creER*^*T2*^ mice). To elicit cre-mediated excision of this stop codon, we injected mice intraperitoneally with 100 mg tamoxifen (Sigma, T5648)/kg mouse/day for 8 days, preceding any experimental manipulations. Same-litter cre recombinase-negative control mice (*Ucp2KI*^*fl/fl*^) were also injected with tamoxifen to control for any potential biological impacts of tamoxifen.

Rosiglitazone (RSG) was fed to WT mice by grinding 4 mg pills (Avandia, GSK) with a mortar and pestle and mixing them in to ground normal mouse chow. We measured daily food consumption and adjusted the amount of RSG used based on food consumption. RSG was fed to mice beginning 2 days prior to microbead injection and does not alter intra-ocular pressure. During this study, we estimate an average RSG consumption of 28.2 mg RSG/kg mouse/day.

### Microbead Injection

We modeled glaucoma in mice by elevating IOP. We increased IOP in 2-4 month old mice of both genders as previously described (Cone et al., 2012). At least 24 hours prior to bead injection, we took a baseline IOP measurement Prior to bead injection, IOP is stable and is well represented by a single measurement. Immediately prior to bead injection we anesthetized mouse corneas topically with proparacaine hydrochloride (0.5%) eyedrops and systemically with an intraperitoneal injection of 100 mg/kg Ketamine/10 mg/kg Xylazine. While anesthetized, we injected 6 µm (2 µL at 3×10^6^ beads/µL; Polysciences, Cat#: 07312-5) and 1 µm (2 µL at 1.5×10^7^ beads/µL; Polysciences, Cat#: 07310-15) polystyrene microbeads through a 50-100 µm cannula in the cornea formed by a beveled glass micropipette connected by polyethylene tubing to a Hamilton syringe (Hamilton Company Reno, NV, USA). As an internal control, 4 µL of sterile 1x phosphate buffered saline (PBS) was injected in to the contralateral eye. We measured postoperative IOP every 3 days for 30 days. Following terminal IOP measurements, mice were asphyxiated using a Euthanex SmartBox system, which automatically controls CO_2_ dispersion, followed by cervical dislocation.

### IOP Measurement

Intra-ocular pressure (IOP) was measured in mice anesthetized by 1.5% isoflurane in air (v/v) using an Icare^®^ TonoLab (Icare Finland Oy, Espoo, Finland) rebound tonometer, both before and after injection with polystyrene microbeads. Each reported measurement is the average of 18 technical replicates/mouse/eye. Mice were included in this study if their individual IOP was elevated by ≥3 mmHg or if a paired t-test of IOP over time between microbead and PBS-injected eyes was statistically significant (p<0.05). Baseline and bead-injected IOPs were compared between mouse strains to confirm the absence of any genotype-dependent differences in IOP increase.

### Histology and Immunocytochemistry

Immunolabeling of sectioned retinal tissue was performed as previously described (Pinzon-Guzman et al., 2011). Briefly, whole eyes were fixed in 4% paraformaldehyde (Electron Microscopy Sciences, Hatfield, PA, USA) in 1x PBS overnight at 4°C. The next day, eyes were divided in half with a scalpel blade. One half was frozen and sectioned, while the other was labeled as a whole-mount. Frozen tissues were embedded in a 2:1 mixture of 20% sucrose and OCT (Electron Microscopy Sciences), cooled to -20°C, and cut at a 10 µm thickness. Samples for each experiment were located on the same slide to control for assay variability. Prior to immunohistochemical labeling, we unmasked antigens in a pH 6.0 sodium citrate buffer. Subsequent labeling of oxidative protein carbonyls was performed using an OxyIHC kit (EMD-Millipore, Cat#: S7450). Derivatization of protein carbonyl groups and all subsequent steps were performed in accordance with the manufacturers instructions. Staining intensity was derived using the H-DAB vector of the ImageJ Color Deconvolution tool background was subtracted from each image, resulting in a numerical semiquantitative measure of oxidative tissue stress. Tissue was imaged using an Olympus BX50 microscope. In this and all other experiments, the acquisition parameters for any given label were held constant.

Post-fixation, retinal whole mounts were permeabilized with 0.2% Triton-X-100 in PBS, blocked with 5% nonfat milk, and incubated in rabbit anti-RBPMS antibody (EMD Millipore) for 6 days at 4°C. Tissue was incubated in secondary antibody and 1 µg/mL Hochest-33258 overnight at 4°C prion to washing and mounting with 0.5% n-propyl gallate in 1:1 glycerol: PBS. Whole-mount tissue was imaged on a Fluoview FV1000 confocal microscope (Olympus).

### Retinal Ganglion Cell Counting

RGCs were counted in retinal whole-mounts using the marker RBPMS (Rodriguez et al., 2014) across 3-4 317.95 µm x 317.95 µm fields, with each field centered 1000 µm from the optic nerve head. Cell counts were converted to measurements of RGC density, averaged for a single retina, and RGC loss was calculated as the difference in mean RGC density between PBS-injected and contralateral bead-injected retinas. The counter was blinded to the identity of each sample. We did not find a significant effect of retinal quadrant on RGC density normally or with elevated IOP, and our images were therefore taken across all retinal quadrants. The mean±SEM and median RBPMS^+^ RGC densities in PBS-injected retinas (pooled from wild-type C57BL6/J and Ucp2^KI^ controls) were 4758±113 and 4738 cells/mm^2^, respectively. The mean±SEM and median RBPMS^+^ RGC densities in Bead-injected retinas were 3957±152 and 3858 cells/mm^2^, respectively, leading to an average 17% cell loss 30 days following bead injection.

### RNA Isolation and Quantitative Real-Time PCR

Flash frozen cells or tissue were lysed in TRIzol (Thermo-Fischer, Cat#:15596018) and RNA precipitated using the manufacturer’s recommended procedure. Final RNA concentration was measured using a NanoDrop ND-1000 Spectrophotometer prior to reverse transcription. We reverse transcribed 300-1000 µg RNA using SuperScript III (Thermo-Fischer, Cat#: 18080093) with random hexamers. cDNA was amplified with iQ SYBR Green Supermix (Bio-Rad, Cat#: 1708882) and amplified on a Bio-Rad iCycler. *Ucp2* primer sequences were F: 5’— GCT CAG AGC ATG CAG GCA TCG—3’ and R: 5’— CGT GCA ATG GTC TTG TAG GCT TCG —3’. TATA-box binding protein (*Tbp)* primer sequences were F: 5’—ACC TTA TGC TCA GGG CTT GGC C—3 R: 5’—GTC CTG TGC CGT AAG GCA TCA TTG—3’. Cq’s from *Ucp2* amplification were normalized against *Tbp* and controls using the ΔΔC_t_ method.

### Primary Astrocyte Culture

Primary mouse cortical astrocytes were isolated from postnatal day 1-4 mice as previously described (Sarafian et al., 2010; Lapp et al., 2014). Briefly, mice were decapitated and brains were removed from the skull. In tissue culture medium, a ∼1 cm portion of superior cerebral cortex was pinched off of the brain using curved forceps. Meninges were removed, and the tissue was triturated with a sterile flame-polished glass Pasteur pipette until it formed a single cell suspension, approximately 20x. The suspension was filtered through a 70 µm cell strainer (Corning, Cat#: 352350) to remove larger debris, centrifuged at 500 x g and 4°C for 5 min, resuspended in growth medium (Dulbecco’s Modified Eagle’s Medium/Ham’s F12 supplemented with 2 mM L-glutamine, 15 mM HEPES, 10% fetal bovine serum, and 10 ng/mL gentamicin), and plated in a T-25 tissue culture flask. Cells were grown at 37°C in a 5% CO_2_/balance air atmosphere. After the cells reached confluence, between 7-14 days in vitro (DIV), contaminating cells were shaken off by rotating at 250 RPM overnight. Astrocyte-enriched cultures were plated at 30,000 cells/well on black tissue-culture-treated 96-well plates (Corning, Cat#3603) and used at passage #2 or 3, allowing at least 48 hours following medium replacement before experimentation. All cells used in this study were exposed to 1 µM 4-hydroxytamoxifen (Sigma, Cat#: H6278) for 24 hours prior to studies of *Ucp2* function.

### Measurement of Mitochondrial Membrane Potential and Oxidative Status

We determined mitochondrial membrane potential (Ψ_m_) and oxidative status of primary cortical astrocytes using the mitochondrial membrane potential-sensitive dye TMRE (50 nM, ImmunoChemistry, Cat#: 9103) or the mitochondrial superoxide probe MitoSox (5 µM, Thermo-Fischer, Cat#: M36008), which is selectively targeted to mitochondria. Cells were incubated in either dye in prewarmed assay medium (1x PBS supplemented with 1 mM glucose and 2 mM GlutaMax, Thermo-Fischer, Cat#: 35050-061) for 30 min at 37°C, followed by two washes and imaging. MitoSox fluorescence intensity was measured using the kinetic mode of a microplate reader (BioTek Synergy II), which took serial measurements of MitoSox fluorescence over time. The rate of increase in fluorescence (ΔF) over 10 minutes was divided by initial fluorescent intensity (F_0_) for each well. This rate of increase was normalized to the mean ΔF/F_0_ of control cells. We verified the utility of TMRE as an indicator of Ψ_m_ by simultaneously treating cells with the membrane permeant protonophore Carbonyl cyanide-*4*-(trifluoromethoxy) phenyl-hydrazone (FCCP, 10 µM, Caymen Chemical, Cat#: 15218), which depolarizes Ψ_m_. Similarly, we used the mitochondrial complex III inhibitor Antimycin A (5 µM) to stimulate ROS production and confirm the utility of MitoSox as an indicator of ROS.

### Statistical analysis

Quantified data are represented by that group’s mean±sem unless otherwise indicated. We performed all statistical analyses in GraphPad Prism. We determined the statistical effect of one independent variable on 2 groups using a Student’s t-test or paired sample t-test in cases where samples were matched (e.g. the control was the contralateral eye of the same animal). We analyzed the effect of one variable on >2 groups using a one-way ANOVA with a Bonferroni post-hoc analysis. We analyzed the effect of 2 variables using a 2-way ANOVA with a Bonferroni’s post-hoc analysis. The statistical significance threshold was set at p<0.05 for all tests.

## Results

### Exogenous Uncoupling Agents Decrease the Generation of Mitochondrial ROS

The positive association between mitochondrial membrane potential (Ψ_m_) and the production of reactive oxygen species (ROS) has been well characterized in isolated mitochondria, and we tested the hypothesis that mild mitochondrial uncoupling stimulated by an exogenous protonophore will decrease mitochondrial ROS in intact cells. We treated primary cortical astrocytes with FCCP at a low concentration to uncouple mitochondria without completely dissipating the Ψ_m_, and found that 10 nM FCCP depolarized the mitochondrial membrane potential (Ψ_m_) to 88±4% of control levels, whereas Ψ_m_ was 34±2% of control in astrocytes treated with 10 µM FCCP, a concentration routinely used to maximally depolarize mitochondria (10 µM; Fig. 1A). 10 nM FCCP added to MitoSox loaded cells did not significantly alter mitochondrial superoxide generation (data not shown), so we tested the hypothesis that uncoupling will reduce ROS production by dysfunctional mitochondria. To test this hypothesis we loaded cells with the mitochondrion-targeted superoxide probe MitoSox and treated them with the mitochondrial complex III inhibitor Antimycin A (AA; 5 µM). AA significantly increased the rate of MitoSox oxidation (p<0.001, Fig. 1B), but this increase was partially attenuated in cells simultaneously treated with AA and 10 nM FCCP (p<0.05, Fig. 1B). These data show that uncoupling decreases the generation of ROS by cultured astrocytes with dysfunctional mitochondria.

**Figure 1.**
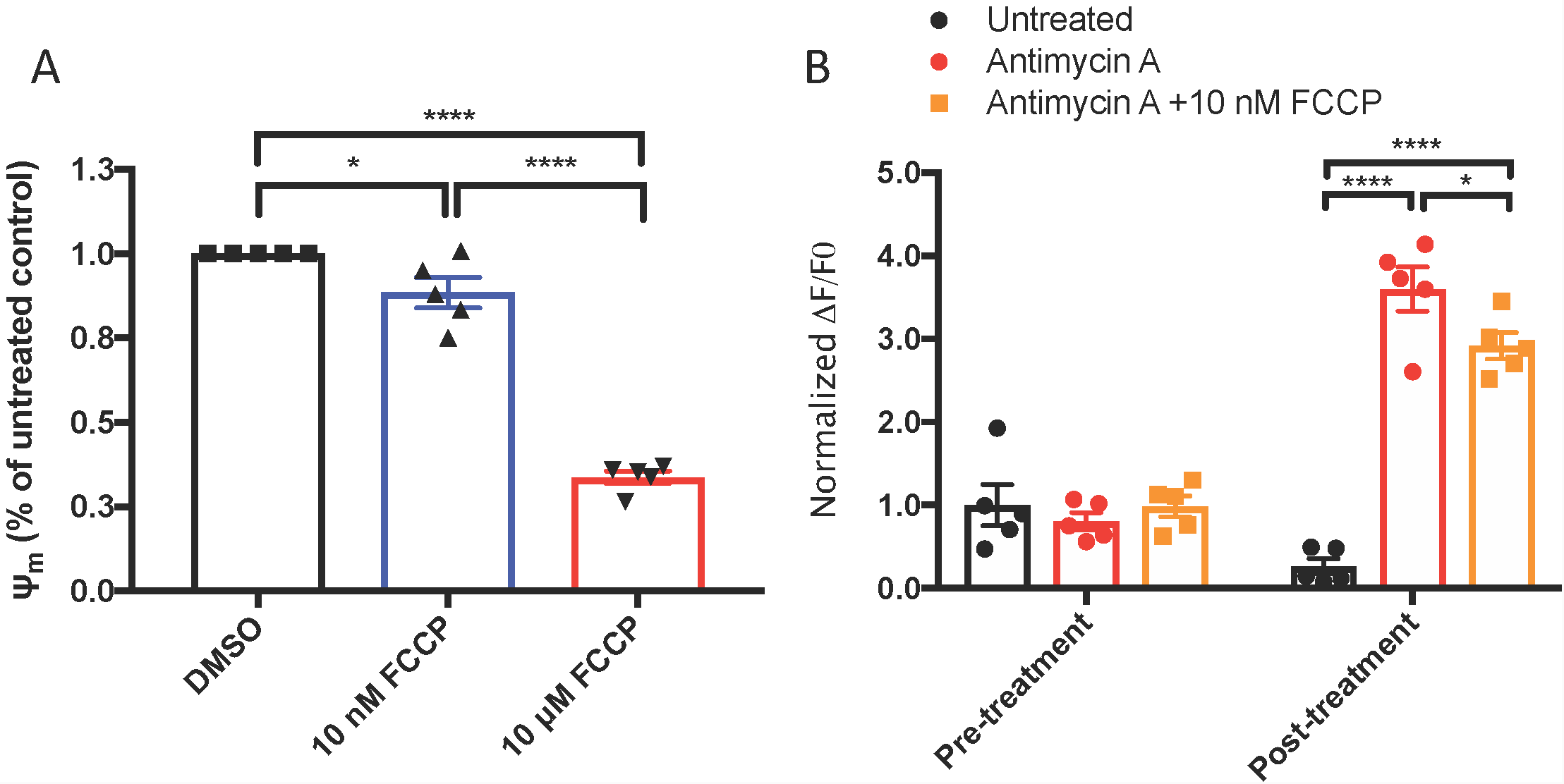
Mitochondrial Uncoupling Decreases Reactive Oxygen Species Production. (**A**) Measurements of TMRE fluorescence as a proxy for mitochondrial membrane potential (Ψ_m_) in cells exposed to vehicle, 10 nM FCCP, or 10 µM FCCP (n=3). (**B**) Increase in MitoSox oxidation over a 5 minute period prior to and following exposure of primary cortical astrocytes to nothing (black circles), 5 uM antimycin A (red circles), or 5 uM antimycin A and 10 nM FCCP (orange circles, n=5). *p<0.05, ****p<0.0001

### Uncoupling Protein 2 Decreases Ψ_m_ and ROS Production

To determine whether mitochondrial uncoupling proteins have the same cellular effects as chemical protonophores on Ψ_m_ and ROS production, we isolated cortical astrocytes from Ucp2^KI^; *GFAP-creER*^*T2*^ mice. *Ucp2* expression is elevated roughly 3-fold in *GFAP-creER*^*T2*^ expressing cells of these mice following exposure to 4-hydroxytamoxifen (Fig. 2A), and we tested the hypothesis that the addition of transgenic *Ucp2* will decrease Ψ_m_ and the generation of ROS. Our data show that Ucp2 knock-in depolarizes Ψ_m_ to 72±7% of control levels (p=0.0095, Fig. 2B), with 10 µM FCCP decreasing TMRE fluorescence to 54±5% of controls (p=0.0002). Increasing Ucp2 levels decreased the production of ROS, monitored by the change in MitoSox fluorescence over time and normalized to the mean fluorescent intensity of Ucp2^KI^ control samples (p=0.043; Fig. 2C). Together, these data show that increased Ucp2 expression decreases Ψ_m_ and mitochondrial ROS, which may be similar in mechanism to the protective effects promoted by 10 nM FCCP.

**Figure 2.**
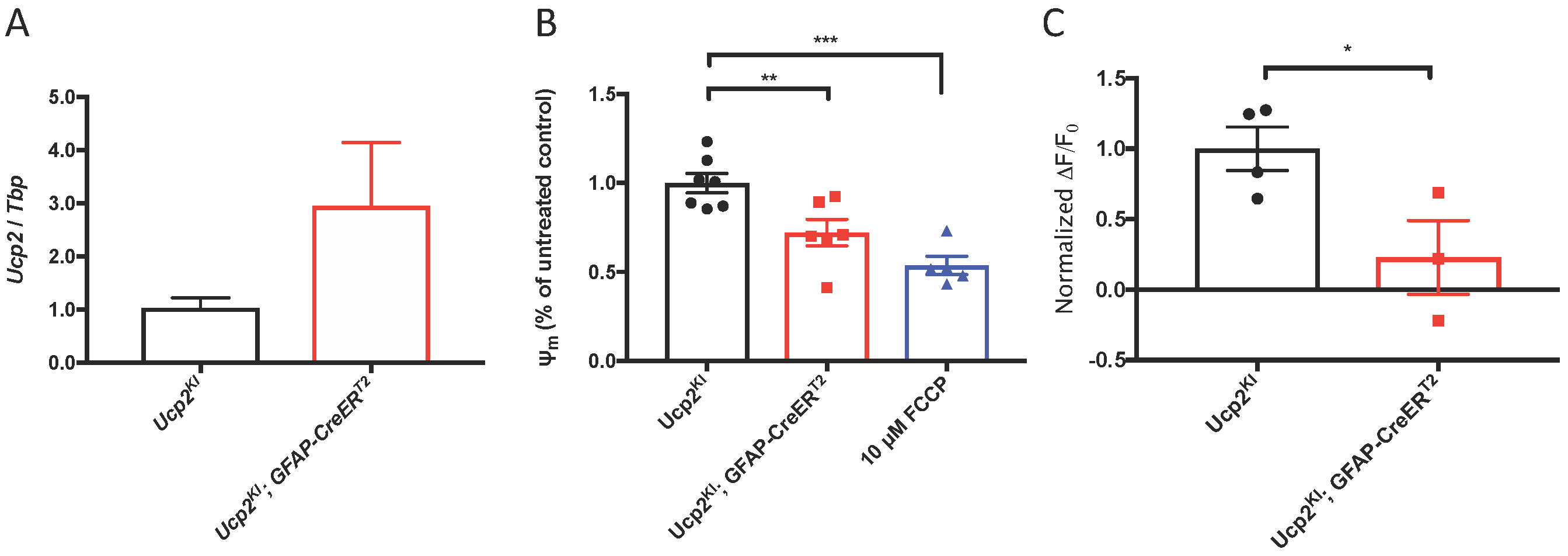
Ucp2 Decreases Ψ_m_, Increasing Respiration and Decreasing Production of ROS. (**A**) Ucp2 gene expression (n=3), (**B**) Relative TMRE fluorescence (Ψ_m_, n=5-7), and (**C**) the relative rate of increase in MitoSox fluorescence (n=3-4) within primary cortical astrocytes isolated from *Ucp2*^*KI*^ and *Ucp2*^*KI*^; *Gfap-creER*^*T2*^ mice. *p<0.05, **p<0.01, ***p<0.005.

### Elevated IOP increases Ucp2 Expression

To determine whether Ucp2 expression is positively or negatively related to the progression of glaucoma, we analyzed publically available data from a microarray (https://www.ncbi.nlm.nih.gov/sites/GDSbrowser?acc=GDS3899) that determined gene expression changes in the retina and optic nerve heads of 10.5 month-old DBA/2J mice and DBA/2J; *Gpnmb*^+^ controls (Howell et al., 2014). DBA/2J are genetically predisposed towards glaucoma, and DBA/2J; *Gpnmb*^+^ controls are genetically identical to these mice, except for in the *Gpnmb* gene, for which these control mice express a wild-type copy. Relative to the housekeeping gene *Tbp, Ucp2* expression is elevated early in glaucoma, but decreases with increasing disease severity (p<0.05, Figure 3A). We confirmed these data in a microbead model of glaucoma, and found that 3 days following microbead-injection, intra-ocular pressure (IOP) is significantly elevated (p<0.05, Fig. 3B). Following IOP measurement, we determined Ucp2 expression in the retinas of these mice, and found that Ucp2 expression increases proportionally with IOP (r^2^=0.8, p=0.0001 Fig. 3C). These data suggest that Ucp2 may play a role in the retinal response to IOP elevation.

**Figure 3.**
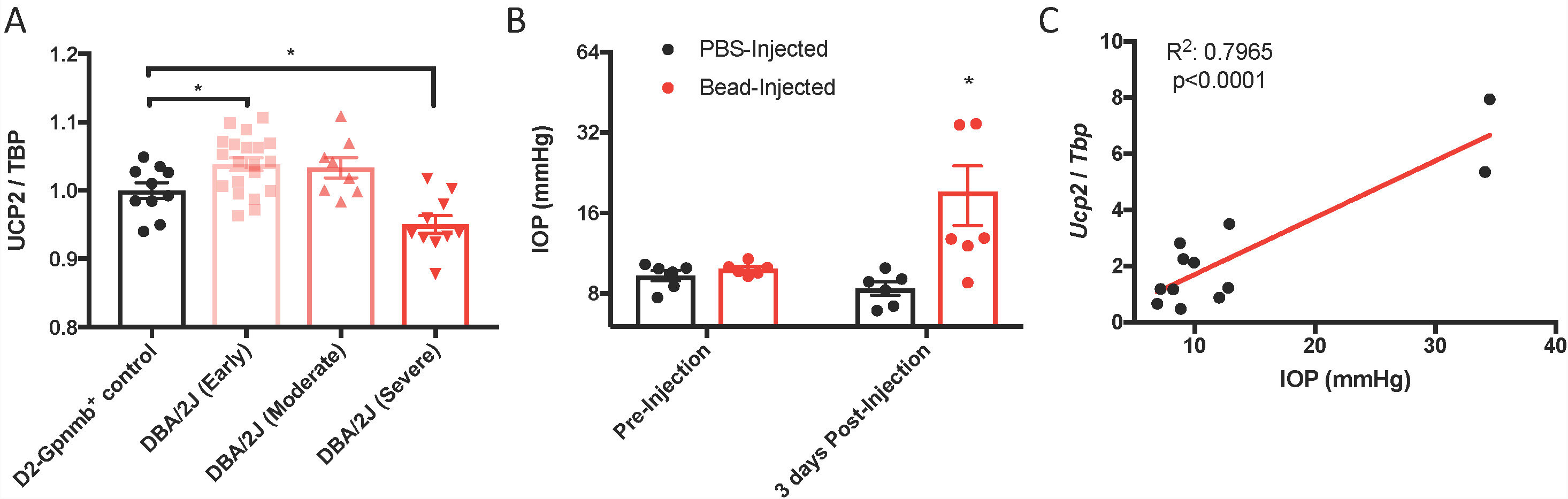
Glaucoma Increases *Ucp2* Expression. (**A**) *Ucp2* expression in a genome-wide microarray of gene expression in the retinas of DBA/2J mice at various stages of glaucoma (n=10-20), and DBA/2J-Gpmnb^+^ controls (n=10) (Howell et al., 2011). (**B**) Intra-ocular pressure (IOP) prior to and following microbead injection (n=6) and (**C**) expression of *Ucp2* in each of the retinas from these mice as a function of IOP, 3 days following bead injection (n=12). *p<0.05.

### Elevated Ucp2 Expression in RGCs but not Astrocytes or Müller Glia is Protective Against Glaucoma

To determine whether the protective effects of *Ucp2* expression in cells translate to the same *in vivo* system, we used mice in which Ucp2 expression can be increased in *Gfap*-or *Thy1*-expressing cells following exposure to tamoxifen (Fig. 4A). Following eight consecutive 100 mg/kg/day injections, we found that *GFAP-creER*^*T2*^ expression increased *Ucp2* transcript levels to 165±14% of control (p<0.01, n=6), and *Thy1-creER*^*T2*^ increased Ucp2 to 229±77% of *Ucp2*^*KI*^ controls (p<0.05, n=3, Fig. 4B). Following exposure to tamoxifen, cells express eGFP, pictured in Fig. 4C, though notably, YFP, which spectrally overlaps with eGFP, is expressed both before and after tamoxifen injection in *Thy1-creER*^*T2*^ retinas (Fig. 4C). The white arrows in Fig. 4C indicate regions of each cre variant that are structurally distinct, including müller glia fibers (top) and retinal ganglion cell soma (bottom).

**Figure 4.**
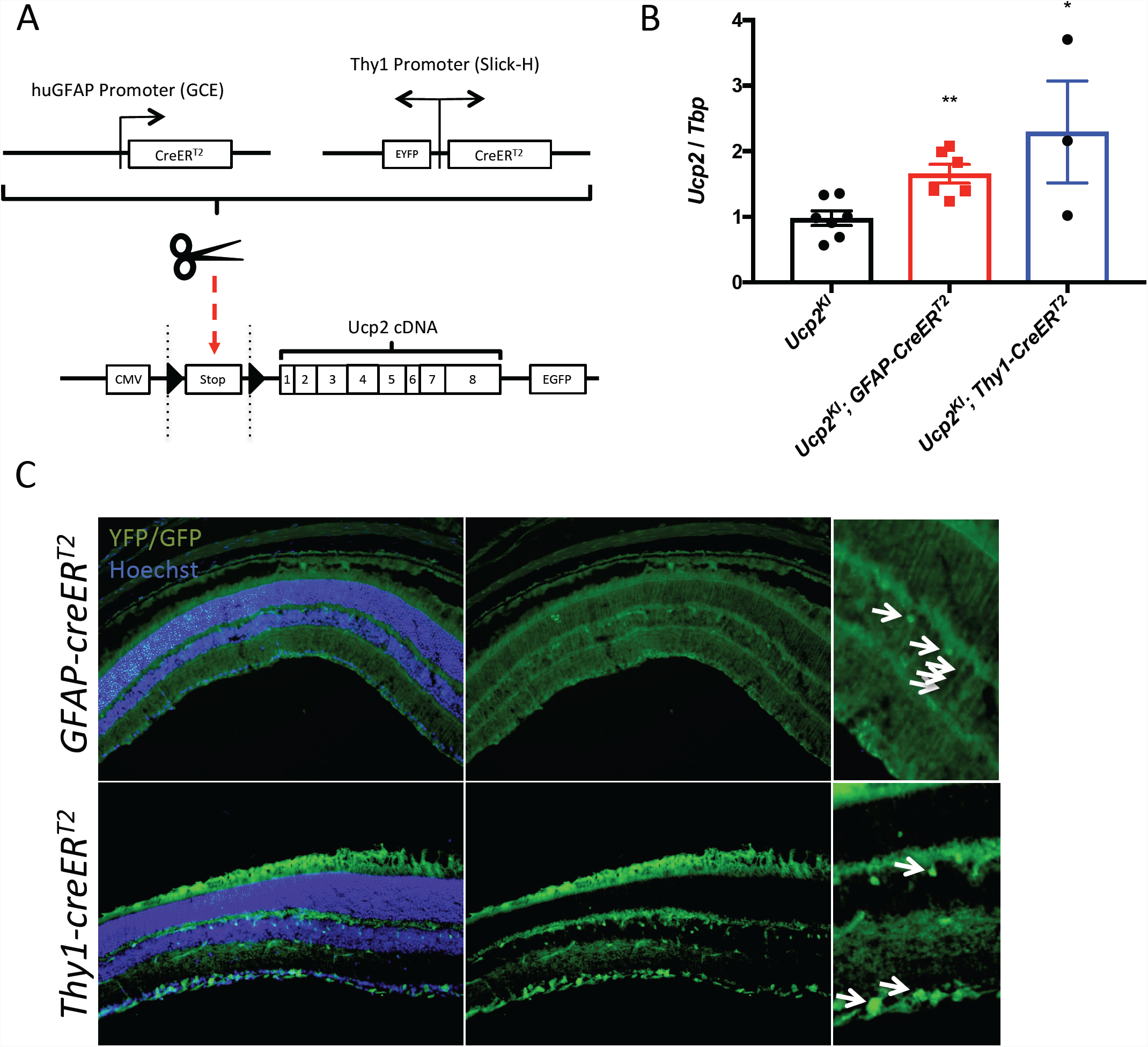
Effects of *Thy1-creER*^*T2*^ and *Gfap-creER*^*T2*^ on Retinal *Cre* Recombinase Localization and Ucp2 expression. (**A**) Gene diagram of transgenic *Ucp2* and *cre* recombinase variants used in the present study. (**B**) Expression of *Ucp2* in control and transgenic mice (n=3). (**C**) Hoechst 33258 (blue)-labeled frozen sections of *Ucp2*^*KI*^; *GFAP-creER*^*T2*^ and *Ucp2*^*KI*^; *Thy1-creER*^*T2*^ retinas, showing the endogenous fluorescence of EGFP and YFP, respectively. White arrows point to Muller glia filaments and cell bodies in *GFAP-creER*^*T2*^ retinas and to RGC soma in *Thy1-creER*^*T2*^ retinas. (**D**) Representative images and quantification of OxyIHC-dependent DAB labeling of fixed *Ucp2*^*KI*^, *Ucp2*^*KI*^; *GFAP-creER*^*T2*^, and *Ucp2*^*KI*^; *Thy1-creER*^*T2*^ retinas, showing that *Ucp2*^*KI*^; *Thy1-creER*^*T2*^ mice generate a significantly fewer oxidative stress-dependent protein carbonyls. *p<0.05, **p<0.05,

We injected microbeads or PBS in to the anterior chambers of these mice, elevating IOP by an average of 5.3 mmHg in *Ucp2*^*KI*^ control mice, 2.4 mmHg in *Ucp2*^*KI*^; *GFAP*-*creER*^*T2*^ mice, and 7.5 mmHg in *Ucp2*^*KI*^; *Thy1*-creER^T2^ mice (Fig. 5A). Bead injection in control mice caused a significant loss in RGCs (1018±88 cells/mm^2^ for a 19±3% reduction in RGCs; n=11) that was attenuated in mice overexpressing Ucp2 in RGCs (*Ucp2*^*KI*^; *Thy1-creER*^*T2*^, 492±175 cells/mm^2^ for a 10±4% reduction; n=9), but not in *Ucp2*^*KI*^; *GFAP*-*creER*^*T2*^ mice (824±225 cells/mm^2^ for a 15±4% reduction, n=6, Fig. 5C-D). These data demonstrate that Ucp2 decreases RGC loss due to elevated IOP over a subacute timeframe, and also that the beneficial effects of Ucp2 are cell autonomous, as Ucp2-overexpression in *GFAP*-positive glia is insufficient to decrease glaucoma-related RGC loss (Fig. 5C). The protective effects of Ucp2 expression coincided with decreases in oxidative protein carbonylation, measured by OxyIHC labeling. Bead-injected retinas from *Ucp2*^*KI*^; *Thy1-creER*^*T2*^ mice (n=4) were labeled 27±6% less strongly than corresponding *Ucp2*^*KI*^ controls (p<0.05, n=7). In contrast, labeling of *Ucp2*^*KI*^; *GFAP*-*creER*^*T2*^retinas was non-significantly reduced by 11±11% (n=3, Fig. 6A-B) relative to controls.

**Figure 5.**
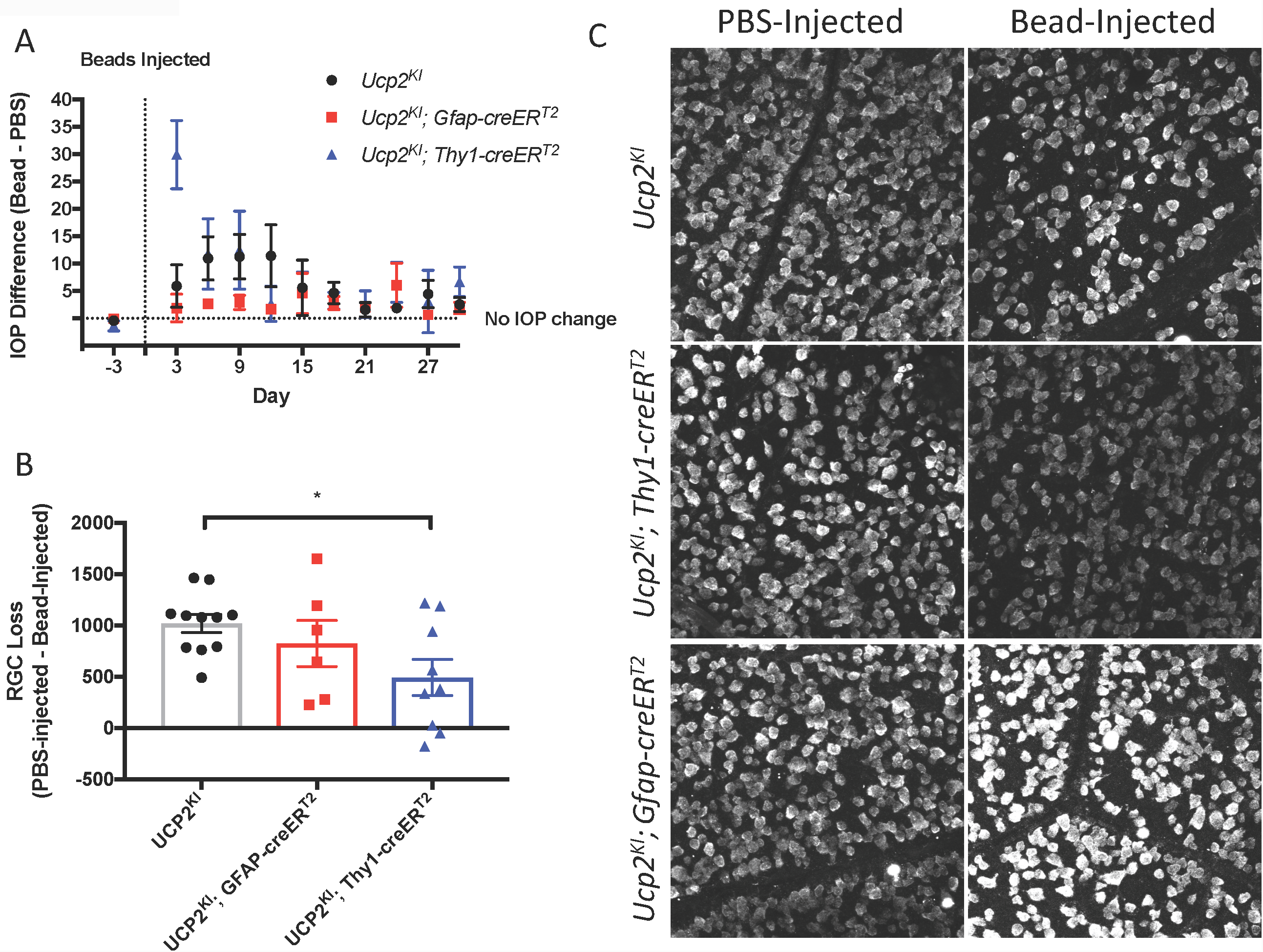
Increased *Ucp2* Expression in RGCs but not Astrocytes and Müller Glia Decreases Retinal Ganglion Cell Loss. (**A**) Difference IOP between bead-and PBS-injected eyes before and following intraocular surgery. Beads injection elevated IOP over a 30 day period in *Ucp2*^*KI*^ (n=11), *Ucp2*^*KI*^; *Gfap-creER*^*T2*^ (n=6), *Ucp2*^*KI*^; *Thy1-creER*^*T2*^ (n=9) eyes. (**B**) RGC loss in retinas in retinal whole-mounts from Bead-and PBS-injected eyes of the indicated genotypes. (**C**) Representative images of retinal whole-mounts labeled for RBPMS, quantified in (**B**). *p<0.05

**Figure 6.**
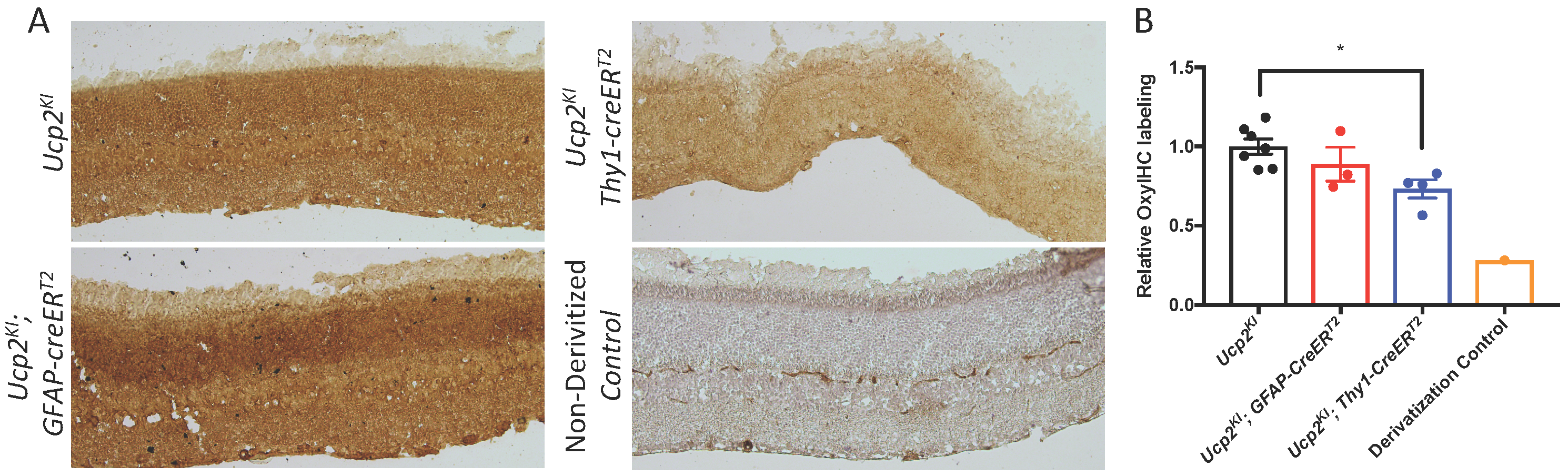
Increased *Ucp2* Expression in RGCs but not Astrocytes and Müller Glia Decreases Oxidative Stress. **(A)** Representative labeling of fixed retinas from bead-injected *Ucp2*^*KI*^ (n=7), *Ucp2*^*KI*^; *Gfap-creER*^*T2*^ (n=3), and *Ucp2*^*KI*^; *Thy1-creER*^*T2*^ (n=4) eyes for oxidative stress-derived protein carbonylation. A single section was labeled without using the 2’,4’-dinitrophenylhydrazine derivitization reagent to confirm the specificity of the primary antibody to towards dinitrophenylhydrazones. The intensity of DAB labeling is quantified in (**B**).

### Transcriptional Activation of Ucp2 is Insufficient to Decrease Microbead-Induced RGC Loss

Past literature suggests that Ucp2 transcription is in part regulated by a PGC1-α/PPAR-γ axis (Chen et al., 2006; Donadelli et al., 2014). Rosiglitazone (RSG) is an FDA-approved PPAR-γ agonist. We confirmed that retinal *Ucp2* expression can increase 24 hours following exposure to 10 mg/kg rosiglitazone (Fig. 7B), and hypothesized that due to transcriptional activation of Ucp2, dietary rosiglitazone confers the same resistance to damage in glaucoma as transgenic Ucp2 overexpression. To test this hypothesis, we increased IOP in control-and RSG-fed WT mice (Fig. 7A) and measured RGC loss 30 days following bead injection. The difference in RGC density between the PBS-and bead-injected eyes in WT control mice was 569±170 cells/mm^2^ (n=3), compared to 961±210 cells/mm^2^ in RSG-fed mice (n=4). The degree of cell loss was generally lower than in Ucp2^KI^ controls, which can be explained by the more advanced age of these mice (3.8 months), which has been demonstrated to reduce the effectiveness of RGC loss following bead injection (Cone et al., 2010). Regardless, the result of this pilot study on the effects of RSG ran contrary to our expectations and did not decrease retinal ganglion cell death, and in fact appeared to non-significantly increase RGC loss (Fig. 7C-D).

**Figure 7.**
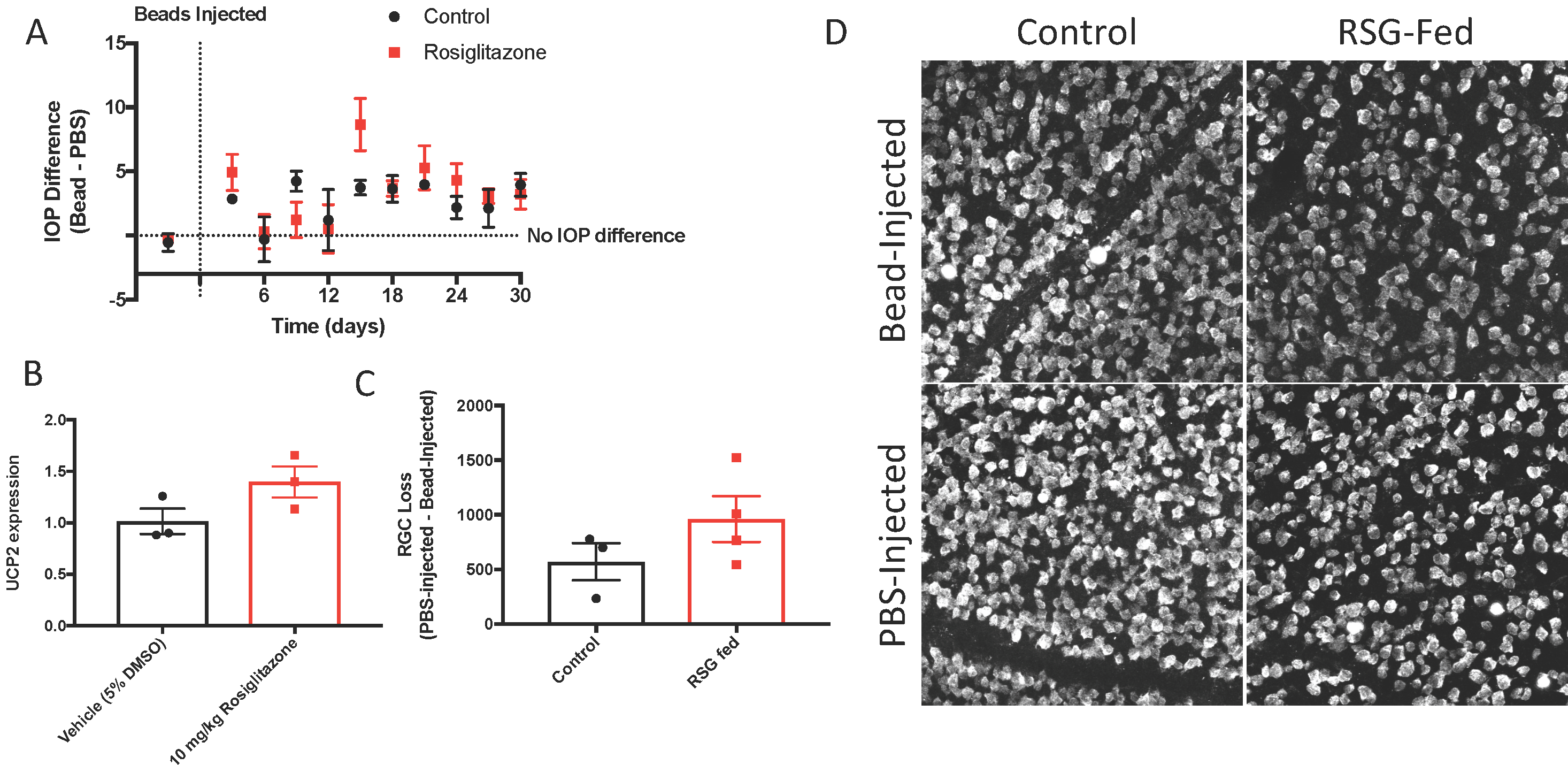
Rosiglitazone Increases *Ucp2* transcription but does not Alter Glaucomatous RGC loss. (**A**) Microbead injection increases IOP to a similar extent in Control-(n=3) and rosiglitazone (RSG)-fed mice (n=4). (**B**) Intraperitoneal RSG increases Ucp2 transcript levels in the mouse retina (n=3) (**C**) increased IOP leads to RGC loss, though dietary RSG does not appear to alter RGC loss. (**D**) Representative RBPMS labeling in whole-mount retinas from PBS-and Bead-injected WT mice.

## Discussion

Mild, homeostatic levels of ROS are important signals of mitochondrial damage (Frank et al., 2012) among other physiological signals (Angelova and Abramov, 2016). When ROS production exceeds the capacity for detoxification by antioxidants, they damage cellular components in a variety of pathogenic conditions. In glaucoma, mitochondria are damaged (Abu-Amero et al., 2006), suggesting that the retina is oxidatively compromised (Feilchenfeld et al., 2008; Chidlow et al., 2017).

Partial dissipation of mitochondrial Ψ_m_ may decrease the generation of mitochondrial ROS during glaucoma (Fig. 6). Using a cell culture system, we found that decreases in Ψ_m_ also decrease ROS production following mitochondrial dysfunction (Fig. 1), similar to data generated using isolated mitochondria (Korshunov et al., 1997; Miwa et al., 2003). We also confirmed that as with other studies (Diano et al., 2003; Lapp et al., 2014), expression of the mitochondrial uncoupling protein *Ucp2* is negatively correlated with both Ψ_m_ and mitochondrial ROS (Fig. 2,6). *Ucp2* expression is altered during different stages of glaucoma, and appears to increase with increasing IOP (Fig. 3). However, we note that it is likely that during later stages of glaucoma, *Ucp2* levels may fall, as we have seen in the data from (Howell et al., 2011). We believe that the decrease in *Ucp2* later in the disease may be a reflection of the decrease in ganglion cell number, which decreases with increased glaucoma severity. If increased ganglion cell *Ucp2* expression reflects a physiological response to increased ROS in glaucoma, artificially increasing Ucp2 may increase the ability of that stress response to increase cell survival. We indeed found that following microbead-injection, we see an increase in retinal *Ucp2* expression (Fig. 5). The hypothesis that *Ucp2* improves cell survival following a cellular stressor is also strongly supported by previous studies (Diano et al., 2003; Mattiasson et al., 2003; Andrews et al., 2005; Barnstable et al., 2016), but the novelty of our study is that in rodent models of glaucoma, retinal ganglion cell death is progressive over time (Huang et al., 2018), suggesting that Ucp2 is not exclusively protective during acutely stressful conditions, but also during chronic neurodegeneration, decreasing the accumulation of oxidative damage (Fig. 6) and bead-induced retinal ganglion cell loss.

Ucp2-mediated neuroprotection is dependent on cell type, as we show that increased *Ucp2* levels in *Gfap*-expressing glia does not significantly alter retinal ganglion cell loss or oxidative stress-derive protein carbonyls compared to controls (Fig. 5-6). A much larger sample size may be sufficient to demonstrate a neuroprotective effect, supported by a trend towards decreased oxidative stress and RGC loss in *Ucp2*^*KI*^; *GFAP*-*creER*^*T2*^ mice, but overall the data argue for much weaker if any *Ucp2*-mediated neuroprotection from glial cells than from RGCs. This seems to suggest that changes in mitochondrial dynamics within *Gfap*-expressing glia of the retina may not be central for the progression of glaucoma, which is unexpected given the many changes they undergo over the course of the disease (Woldemussie et al., 2004) and the protection they give to RGCs (Kawasaki et al., 2000).

Rosiglitazone is a PPAR-γ-dependent transcriptional activator of *Ucp2* (Medvedev et al., 2001; Chen et al., 2006), and increases retinal *Ucp2* expression (Fig. 7), but does not seem to promote *Ucp2* mediated neuroprotection in the microbead model of glaucoma. PPAR-γ appears to be expressed with high specificity in müller glia cells of the rodent retina (Zhu et al., 2013), and while the failure of rosiglitazone to protect RGCs was initially surprising, it likely increases the transcription of glial *Ucp2*. *Ucp2* overexpression in *Gfap*-expressing glia failed to protect RGCs, so our experiments using RSG-fed and *Ucp2*^*KI*^; *GFAP*-*creER*^*T2*^ mice largely support each other. PPAR-γ agonism with pioglitazone is sufficient to decrease RGC loss following optic nerve crush in rats (Zhu et al., 2013) or retinal ischemia/reperfusion injury in mice (Zhang et al., 2017), suggesting that glial *Ucp2* expression may be protective against more acute retinal insults. This protection may also result from the increase in neural PPAR-γ following optic nerve crush (Zhu et al., 2013), which would allow for agonist-stimulated Ucp2 mRNA expression that is likely subject to multiple endogenous regulatory mechanisms (Donadelli et al., 2014), unlike the *Ucp2* derived from our transgenic mice (Toda et al., 2016). Overall, a larger study of *Ucp2* and PPAR-γ in retinal disease that uses multiple models and agonists/antagonists may yield a clearer picture that captures the cell type specific dynamics of these factors, and changes during different paradigms of retinal damage.

Overall, our data suggests that the greatest protection against RGC loss can be provided by stimulating Ucp2 expression in RGCs. Expression of this gene in other cell types may not be harmful, but our results suggest that the choice of therapeutic target should be dictated in part by cell type. The expression and activity of this protein is also tightly regulated, so the future studies on *Ucp2*-mediated neuroprotection should also focus on the factors that manipulate *Ucp2* transcription, translation, and functional activity.

## Acknowledgments

The authors thank Sabrina Diano, PhD for generously donating the Ucp2KOKI^fl/fl^ mice that were the progenitors of mice used in this study. We also thank Evgenya Popova, PhD for her thorough discussions on and critical review of this research.

## Author Contributions Statement

DTH performed the experiments, analyzed them, and wrote the first draft of the manuscript. Both DTH and CJB conceived of the study topic and design, as well as revised the submitted manuscript.

